# The association between Hfq and RNase E in long-term nitrogen starved *Escherichia coli*

**DOI:** 10.1101/2021.04.19.440462

**Authors:** Josh McQuail, Agamemnon J. Carpousis, Sivaramesh Wigneshweraraj

**Affiliations:** MRC Centre for Molecular Bacteriology, Imperial College London, London, SW7 2AZ, UK; Laboratoire de Microbiologie et de Génétique Moléculaires (LMGM), Centre de Biologie Intégrative (CBI), Université de Toulouse, CNRS, UPS, Toulouse, France

## Abstract

Under conditions of nutrient adversity, bacteria adjust metabolism to minimise cellular energy usage. This is often achieved by controlling the synthesis and degradation of RNA. In *Escherichia coli*, RNase E is the central enzyme involved in RNA degradation and serves as a scaffold for the assembly of the multiprotein complex known as the RNA degradosome. The activity of RNase E against specific mRNAs can also be regulated by the action of small RNAs (sRNA). In this case, the ubiquitous bacterial chaperone Hfq bound to sRNAs can interact with the RNA degradosome for the sRNA guided degradation of target RNAs. The RNA degradosome and Hfq have never been visualised together in live bacteria. We now show that in long-term nitrogen starved *E. coli*, both RNase E and Hfq co-localise in a single, large focus. This subcellular assembly, which we refer to as the H-body, forms by a liquid-liquid phase separation type mechanism and includes components of the RNA degradosome, namely, the helicase RhlB and the exoribonuclease polynucleotide phosphorylase. The results support the existence of an hitherto unreported subcellular compartmentalisation of a process(s) associated with RNA management in stressed bacteria.

## 1. Introduction

The synthesis and degradation of mRNA underpins the ability of bacteria to adapt the flow of genetic information to efficiently respond to specific environmental stimuli. In addition, non-coding small RNAs (sRNAs) contribute to the regulation of bacterial gene expression by acting as multitarget repressors and activators of gene expression by binding to the 5’ untranslated regions of mRNAs. In *Escherichia coli* and related bacteria, the endoribonuclease RNase E, an essential enzyme, is involved in the degradation of most mRNAs (Ait-Bara & Carpousis, 2015, Bandyra *et al*., 2013, Vargas-Blanco & Shell, 2020) and the biogenesis of sRNAs from the 3’ untranslated regions of mRNAs (Chao *et al*., 2017, Chao & Vogel, 2016, Chao *et al*., 2012). RNase E has a preference for 5’ monophosphorylated RNA substrates and the RNA phosphohydrolase RppH converts 5’ triphosphorylated mRNA into mRNA with 5’ monophosphorylated ends. RNase E can also cleave RNA in a 5’ end independent manner and genome-wide surveys of RNase E cleavage sites have revealed that this represents the major mRNA cleavage pathway in *E. coli* and *Salmonella* (Chao *et al*., 2017, Clarke *et al*., 2014). The catalytic domain of RNase E (amino acid residues 1-510) is located at the N terminus of the protein. The large noncatalytic carboxyl terminal region of RNase E (amino acid residues 511-1061) is intrinsically disordered and serves as the scaffold for the assembly of a multienzyme complex known as the RNA degradosome (Callaghan *et al*., 2005, Callaghan *et al*., 2004). The ‘minimal’ RNA degradosome of *E. coli* consists of the DEAD-box helicase RhlB and the exoribonuclease polynucleotide phosphorylase (PNPase). Both PNPase and RNase E carry out RNA degradation and RhlB facilitates this by unwinding stem-loops within the RNA targets (Ait-Bara & Carpousis, 2015, Bandyra *et al*., 2013, Vargas-Blanco & Shell, 2020). The glycolytic enzyme enolase is an auxiliary component of the RNA degradosome and also interacts with carboxyl terminal region of RNase E; it is suspected to have a regulatory role in mRNA degradation in response to phosphosugar (Chandran & Luisi, 2006) and anaerobic stress (Murashko & Lin-Chao, 2017). For the sake of simplicity, we will refer to the RNA degradosome containing RNase E, RhlB and PNPase as the minimal RNA degradosome complex.

The carboxyl terminal region of RNase E contains a short amphipathic α-helix, called the membrane targeting sequence (MTS; amino acid residues 565-585), which allows it to interact with the inner cytoplasmic membrane of *E. coli* (Taghbalout & Rothfield, 2007, Taghbalout *et al*., 2014, Khemici *et al*., 2008). Strahl et al showed that RNase E diffuses freely over the entire surface of the inner cytoplasmic membrane in exponentially growing *E. coli* cells but occasionally forms multiple short-lived puncta (Strahl *et al*., 2015). This led the authors to suggest that the RNase E puncta either represented the RNA degradosome or a stable complex of the RNA degradosome with polysomes. Recently, Al-Husini et al showed that the α-proteobacterial RNase E of *Caulobacter crescentus*, which lacks an MTS, is located in the interior of the cell and assembles into liquid-liquid phase separated, short-lived clusters of 0.2-0.4 µm in diameter. These clusters, which are called BR-bodies, recruit RNA degradosome components (in this case RhlB and aconitase, which is the equivalent of enolase) particularly in stressed *C. crescentus* cells (Al-Husini *et al*., 2018). In a subsequent study, Al-Husini et al showed that BR-bodies were specifically enriched in poorly translated mRNAs and sRNAs and functioned to promote efficient, most likely sRNA-guided (see below), degradation of mRNAs in stressed *C. crescentus* cells (Al-Husini *et al*., 2020).

The ubiquitous bacterial chaperone Hfq, when bound to sRNAs, can interact with RNase E for the sRNA-guided degradation of target mRNAs. This RNase E-Hfq/sRNA complex has been proposed to be distinct from the canonical minimal RNA degradosome (Morita *et al*., 2005). However, a study by Bruce et al used structural and biophysical approaches to provide evidence that Hfq/sRNA can interact with the RNA degradosome (Bruce *et al*., 2018). The importance of the sRNA guided association of Hfq with RNase E for targeted RNA degradation in *E. coli* and related bacteria is undisputed (Mackie, 2013, Vargas-Blanco & Shell, 2020, Morita & Aiba, 2011, Ikeda *et al*., 2011, Morita *et al*., 2005). It is not known whether Hfq is part of BR-bodies described in *C. crescentus* (Al-Husini *et al*., 2018). Recently, we demonstrated that Hfq forms single, large (∼0.3 µm), long-lived foci in long-term nitrogen (N) starved *E. coli* (McQuail *et al*., 2020). In this study, we explored whether the minimal RNA degradosome is a constituent of the Hfq foci and thereby represent a compartmentalised subcellular assembly involved in RNA degradation in long-term N starved *E. coli*.

## 2. Results

### 2.1 RNase E assembles into a single, large focus in long-term N starved *Escherichia coli*

Previously we used photoactivated localisation microscopy (PALM) to demonstrate that Hfq forms foci in ∼90% of long-term (24 h) N starved *E. coli* cells (McQuail *et al*., 2020). To determine whether RNase E associates with these Hfq foci in long-term N starved *E. coli*, we constructed an *E. coli* strain containing GFP and photoactivatable mCherry (PAmCherry) fused to the carboxyl terminal ends of RNase E and Hfq at their normal chromosomal locations, respectively. Control experiments established that the ability of the Hfq-PAmCherry and RNase E-GFP tagged *E. coli* strain to survive long-term (24 h) N starvation did not differ from that of the untagged parent strain ((McQuail *et al*., 2020) and (Figure 1a)). We initially studied the intracellular distribution of RNase E-GFP in exponentially growing bacteria under N replete conditions (N+), shortly following N run-out in the growth media (N-) and in long-term N starved bacteria following 24 h of N starvation (N-24) by fluorescent microscopy using the experimental setup described by McQuail et al. (McQuail *et al*., 2020). At N+ and N-, the majority of RNase E molecules were evenly distributed in the cell (Figure 1b and 1c). Consistent with previous observations (Strahl *et al*., 2015), we also observed small clusters of RNase E proximal to the inner cytoplasmic membrane (Figure 1b and 1c). However, at N-24, we observed a single, large cluster of RNase E, which we refer to as the RNase E focus, near the pole of ∼46% of cells (Figure 1d). The RNase E foci were similar in size (∼0.3 µm) to the Hfq foci (McQuail *et al*., 2020). Notably, only one RNase E focus was observed within a cell, which was significantly larger and longer-lived (see below) than the multiple RNase E puncta previously observed in exponentially growing *E. coli* cells by Strahl et al. Like the Hfq foci (McQuail *et al*., 2020), the RNase E foci were observed in cells that were starved for N for up to 72 h (see later). Overall, we conclude that RNase E, like Hfq (McQuail *et al*., 2020), forms a single and large focus in long-term N starved *E. coli*.

**Figure 1.**
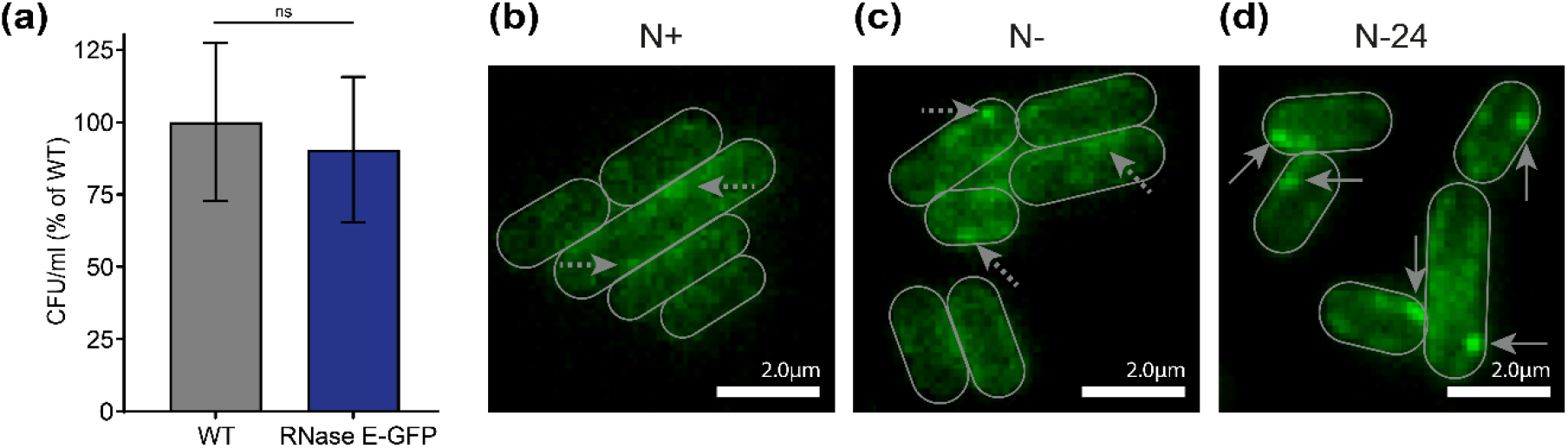
RNase E assembles into a single and large focus in long-term N starved *E. coli* cells. (a) Graph showing the proportion of viable cells in cultures of 24 h N starved WT and GFP tagged RNase E containing *E. coli*, measured by counting CFU and shown as a percentage of WT. Error bars represent standard deviation (n = 3). (b, c, d) Representative fluorescence microscopy images of the distribution of RNase E (GFP-tagged) in *E. coli* cells from (b) N+, (c) N- and (d) N-24. In (b and c), the dashed arrows indicate previously described RNase E puncta. In (d), the solid arrows indicate the RNase E foci.

### 2.2 RNase E and Hfq foci colocalise in long-term N starved *Escherichia coli*

As the RNase E foci (Figure 1) resemble the Hfq foci (McQuail *et al*., 2020) and the functional interaction between Hfq and the RNase E degradosome is undisputed, (Vogel & Luisi, 2011, Chao *et al*., 2017, Bandyra *et al*., 2013, Ait-Bara & Carpousis, 2015), we investigated whether the RNase E and Hfq foci colocalise in long-term N starved *E. coli* cells. At N+ and N-, we only detected (if any at all) minimal overlap between RNase E and Hfq (Figure 2a and 2b). However, at N-24, in all cells that contained a RNase E focus, they colocalised with the Hfq focus (Figure 2c). In other words, within a field-of-view consisting of ∼100-200 cells ∼50% of cells contained an RNase E focus and an Hfq focus that colocalised with it. The colocalisation of the RNase E and Hfq foci persisted for at least 72 h under N starvation (Supplementary Figure 1). Previously we demonstrated that the Hfq foci dispersed when N starvation was alleviated (i.e., upon the addition of N to the *E. coli* cells at N-24) (McQuail *et al*., 2020). We thus considered whether alleviation of N starvation would also cause the concomitant dispersion of the RNase E foci. As shown in Figure 2d, this was indeed the case, and the results suggest that the formation of RNase E and Hfq foci are somehow linked. Consistent with this view, the RNase E foci were undetectable in Δ*hfq E. coli* cells and reappeared when plasmid borne Hfq was supplied to Δ*hfq* bacteria (Figure 2e). Thus, it appears that the formation of the Hfq foci is a requirement for the formation of the RNase E foci. Previously, Morita et al used affinity pull-down experiments to demonstrate that a truncated form of RNase E, lacking the majority of the carboxyl terminal region and thus consisting only of amino acid residues 1-700 (RNase E_1-700_), failed to interact with Hfq (Morita *et al*., 2005). Consistent with this previous result mapping the RNase E region required for interaction with Hfq (Ikeda *et al*., 2011), and supporting the conclusions from results shown in Figure 2e, we failed to detect the RNase E foci in an *E. coli* strain in which RNase E_1-700_ was fused to GFP (Figure 2f). To establish whether Hfq foci formation in the RNase E_1-700_ containing bacteria was affected in any way, we approximated the proportion of Hfq molecules within the foci as a percentage of total number of tracked Hfq molecules within the same field-of-view (consisting of ∼100-200 bacteria) to derive %H_IM_; see McQuail et al for details; (McQuail *et al*., 2020)). As can be seen in Figure 2c and 2f, the %H_IM_ did not differ that much between the WT and RNase E_1-700_ mutant bacteria. In a strain with an RNase E variant lacking the MTS (RNase E_Δ565-585_), RNase E foci formation and its colocalisation with the Hfq foci was also unaffected (∼65% of cells within a field-of-view with ∼100-200 cells contained an RNase E focus and an Hfq focus which colocalised) (Figure 2g) and the %H_IM_ did not differ between the WT and RNase E_Δ565-585_ mutant bacteria. This suggests that RNase E puncta observed previously (which requires inner membrane association (Strahl *et al*., 2015)) and the colocalised RNase E/Hfq foci observed under our conditions are physically distinct. We note that the foci formed by RNase E_Δ565-585_ appear less intense (albeit the same size) than the foci formed by the WT protein. We speculate that this could be due to the intrinsic *in vivo* instability of the RNase E_Δ565-585_ protein (Hadjeras *et al*., 2019). Overall, we conclude that the formation of the large RNase E foci seen in long-term N starved *E. coli* cells, is dependent upon and colocalises with the Hfq foci.

**Figure 2.**
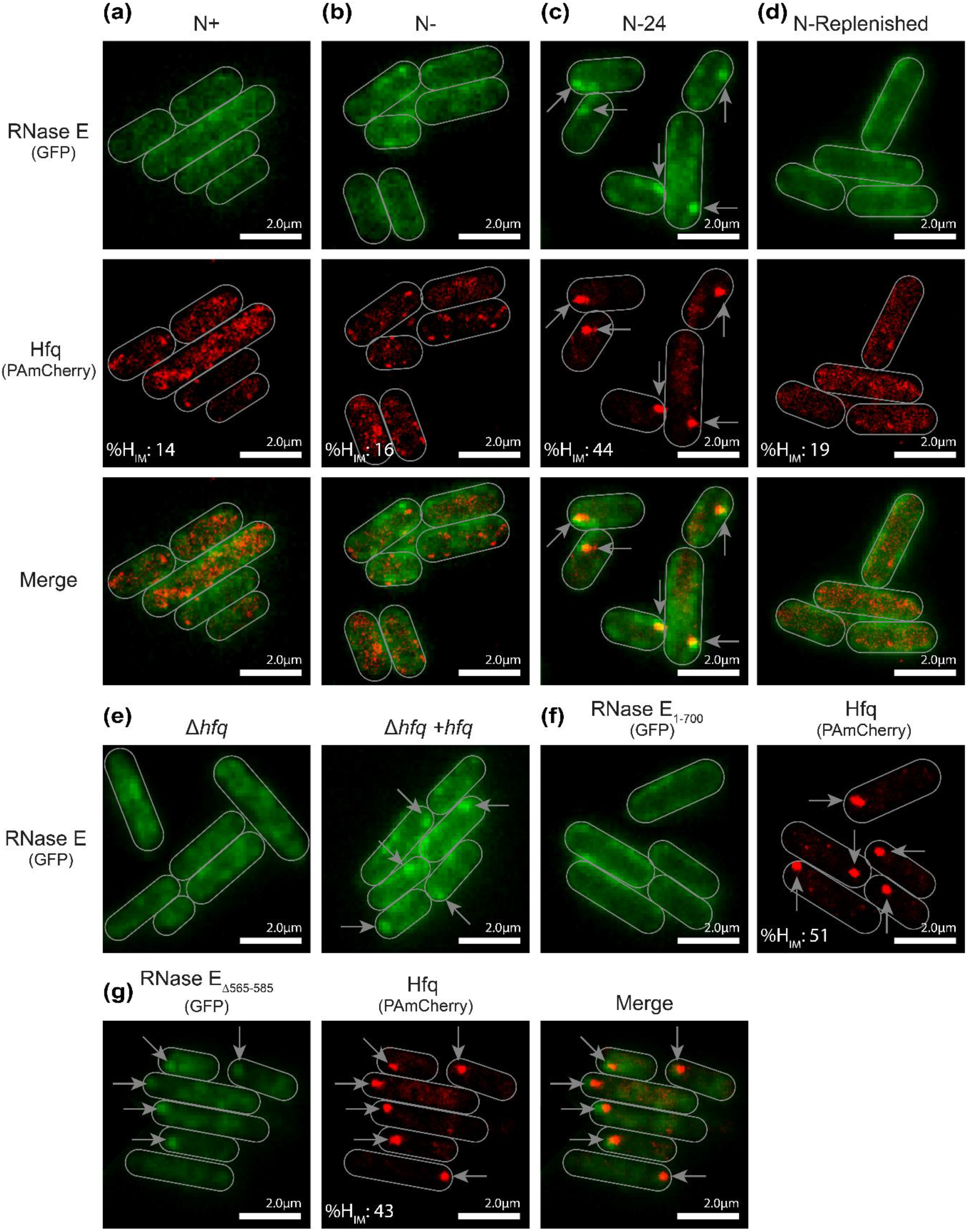
The RNase E and Hfq foci colocalise in long-term N starved *E. coli* cells. (a, b, c, d) Representative fluorescence microscopy (for GFP-tagged RNase E), PALM (for PAmCherry-tagged Hfq) and merged images of RNase E and Hfq in *E. coli* cells, taken at (a) N+, (b) N-, (c) N-24 and (d) 3 h after 24 h N starved bacteria were replenished with N. (e) Representative fluorescence microscopy images of RNase E in 24 h N starved Δ*hfq* and Δ*hfq*+pBAD24-*hfq*-FLAG *E. coli* cells. (f) Representative fluorescence microscopy and PALM images of RNase E and Hfq in 24 h N starved *E. coli* cells containing RNase_1-700._ (g) Representative fluorescence microscopy and PALM images of RNase E_Δ565-585_ and Hfq in 24 h N starved *E. coli* cells. In all figures, the arrows indicate the RNase E and/or Hfq foci. The %H_IM_ values (see text) are indicated in all PALM images.

### 2.3 The colocalisation of RNase E and Hfq foci is dependent on cellular RNA levels in long-term N starved *Escherichia coli*

Recently we reported that, despite running out of N, *E. coli* cells remain transcriptionally active over the initial 24 h under N starvation (Switzer *et al*., 2020), with ∼10% of the transcriptome differentially expressed (242 genes upregulated and 209 genes down regulated). Rifampicin treatment immediately following N run-out had a detrimental effect on cell viability at N-24 (∼50% reduction compared to untreated) suggesting that the transcriptional activity during the initial 24 h following N run-out is important for optimal cell viability (Switzer *et al*., 2020). Rifampicin treatment will lead to the inhibition of *de novo* transcription and subsequent depletion of cellular RNA pool over time. In previous work, we showed that rifampicin treatment immediately following N run-out does not have any signficiant effect on Hfq foci formation at N-24 (McQuail *et al*., 2020). In light of the observation that the Hfq foci and RNase E foci colocalise, we investigated how the treatment of bacteria with rifampicin, immediately following N run-out, affected RNase E foci formation and colocalisation with the Hfq-foci at N-24. As shown in Figure 3, whereas rifampicin, as expected, had no detectable effect on the efficacy of Hfq foci formation (McQuail *et al*., 2020), few, if any, RNase E foci were detected in rifampicin treated bacteria at N-24. The loss of RNase E foci upon rifampicin treatment suggests that cellular RNA levels have a role in RNase E foci formation and that the colocalisation of RNase E and Hfq foci could thus have role in the adaptive response to long-term N starvation.

**Figure 3.**
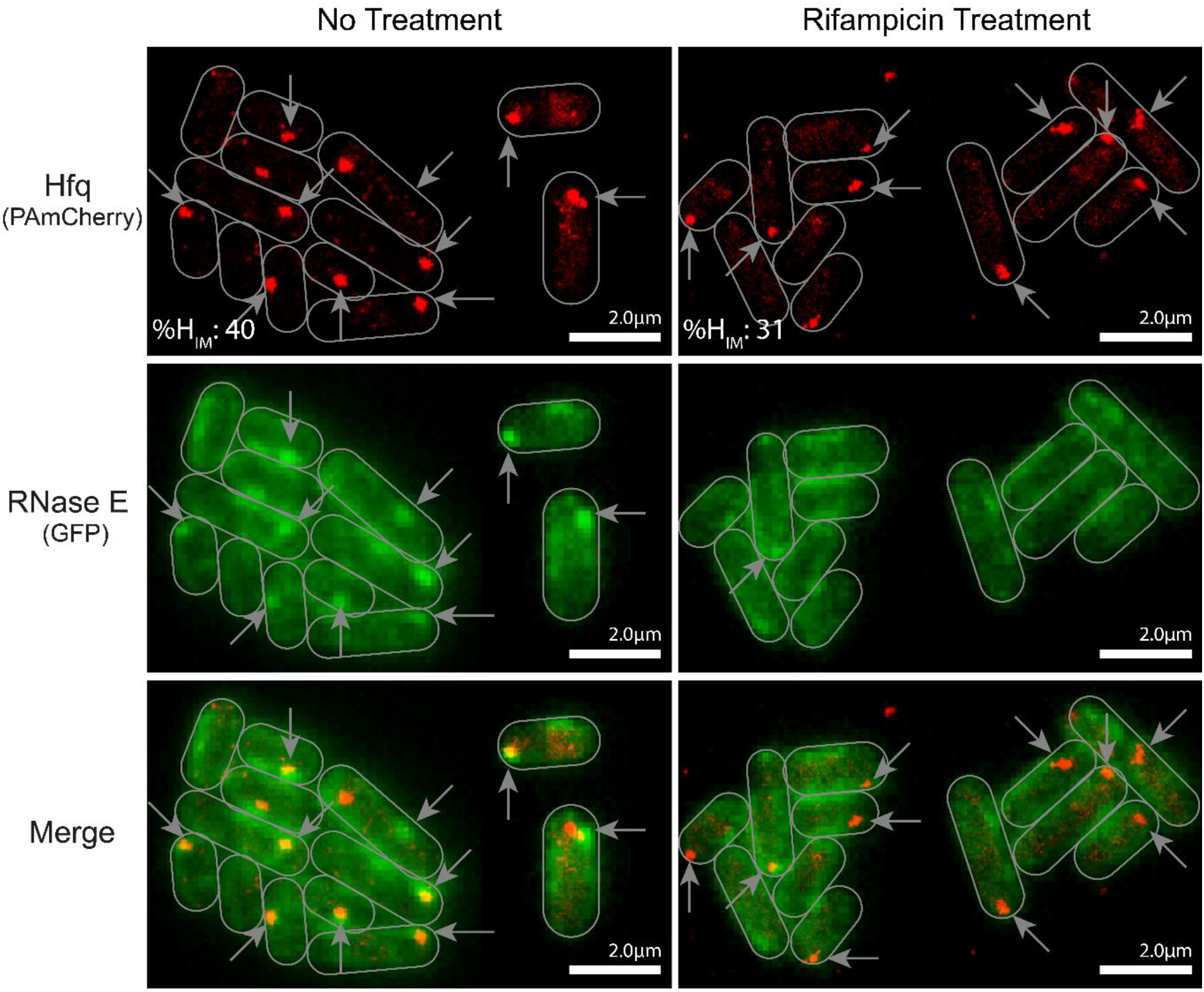
Colocalisation of Hfq and RNase E foci is disrupted by treatment with rifampicin. Representative fluorescence microscopy (for GFP-tagged RNase E) and PALM (for PAmCherry-tagged Hfq) images of the distribution of RNase E and Hfq in untreated and rifampicin (Rif; 100μg/ml) treated (at N-) *E. coli* cells imaged at N-24. The solid arrows indicate the RNase E and/or Hfq foci and the %H_IM_ values are indicated in the PALM images.

### 2.4 The involvement of RhlB and PNPase in the colocalisation of RNase E and Hfq foci in long-term N starved *E. coli* cells

As the minimal RNA degradosome of *E. coli* contains PNPase and RhlB, we investigated whether PNPase and RhlB colocalised with the RNase E/Hfq foci. Control experiments established that the ability of the *E. coli* strains containing PNPase-GFP and RhlB-GFP to survive long-term (24 h) N starvation did not differ from that of the untagged parent strain (Supplementary Figure 2). As shown in Figure 4a and 4b, we observed distinct foci of PNPase and RhlB in *E. coli* cells from N-24 (in ∼57% and ∼45% of cells containing PNPase-GFP and RhlB-GFP, respectively; in both cases within a field-of-view consisting of ∼100-200 cells), which, like the RNase E foci, colocalised with the Hfq foci. We next reasoned that if PNPase and RhlB were indeed part of the RNase E foci (because they interact with RNase E to form the RNA degradosome), then they would not form foci (and thus not colocalise with the Hfq foci) under conditions when the RNase E foci are absent. As the RNase E_1-700_ does not form the foci (and does not contain the interaction sites for PNPase and RhlB) (Figure 2f), we thus investigated whether the RhlB and PNPase can form foci that colocalised with the Hfq foci in an *E. coli* strain containing RNase E_1-700_. As shown in Figure 4c and 4d, respectively, we did not detect any distinct PNPase or RhlB foci in the *E. coli* strain containing the RNase E_1-700_ at N-24; however, as expected (Figure 2f), the formation of the Hfq foci was unaffected. The results thus indicate that the components of the minimal RNA degradosome are recruited by RNase E to the Hfq foci, which specifically form in long-term N starved *E. coli*. We investigated whether the co-localisation of RNase E and Hfq foci in long-term N starved *E. coli* is affected in anyway by the absence of either PNPase or RhlB. As shown in Figure 4e and 4f, RNase E and Hfq foci formation and their colocalisation was unaffected in bacteria devoid of either PNPase or RhlB. Although, we note a slightly higher %H_IM_ in the PNPase mutant bacteria, ∼77% and ∼46% of cells devoid of PNPase and RhlB, respectively, contained colocalised RNase E and Hfq foci. Overall, it seems that a large spatiotemporally separated multiprotein complex, consisting of Hfq, RNase E, RhlB and PNPase, exists in long-term N starved *E. coli*. As the colocalisation of RNase E foci was dependent on the Hfq foci, we have referred to the colocalised feature as the H-body.

**Figure 4.**
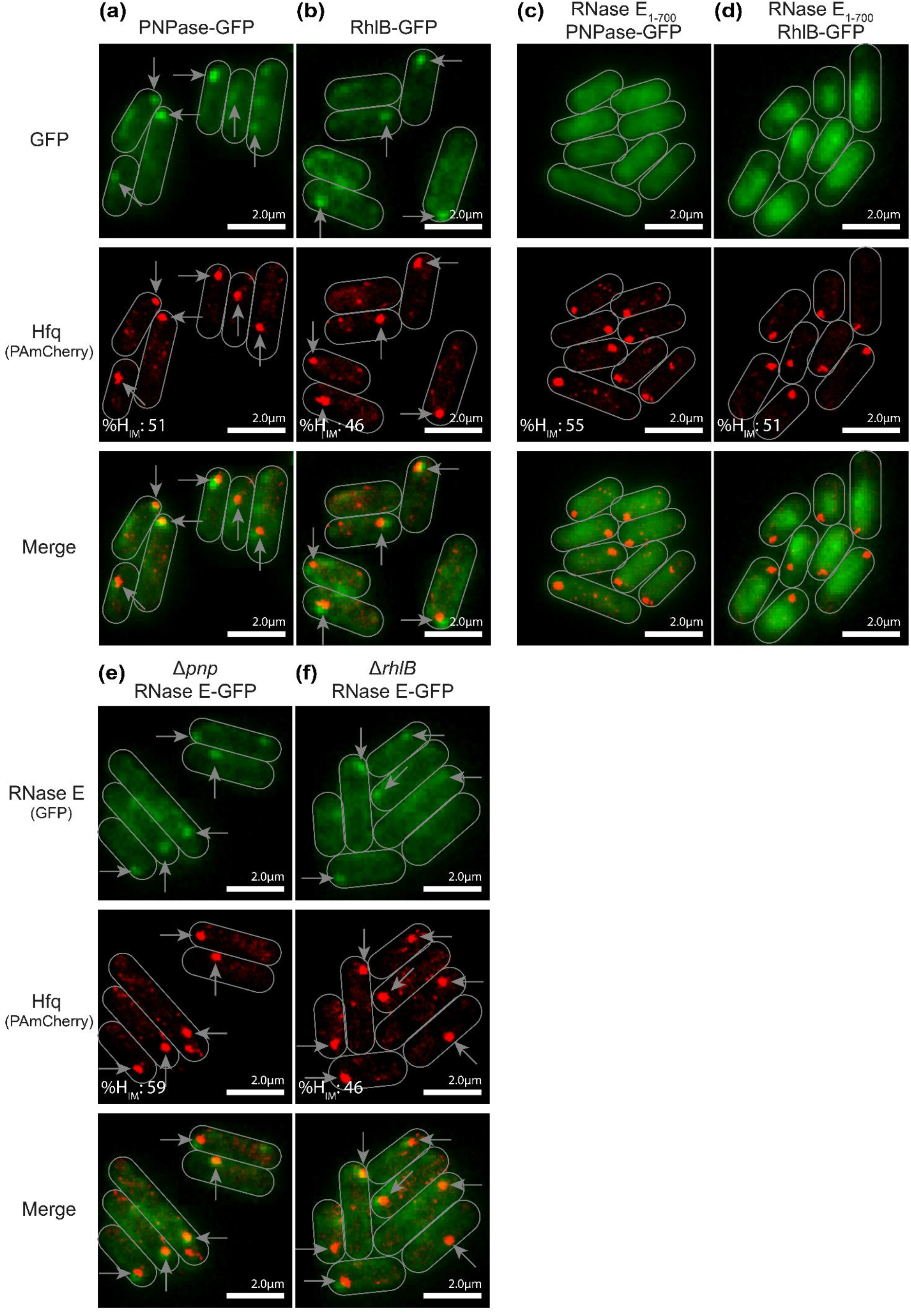
The involvement of RhlB and PNPase in the colocalisation of RNase E and Hfq foci in long-term N starved *E. coli* cells. (a, b) Representative fluorescence microscopy (for GFP-tagged PNPase and RhlB), PALM (for PAmCherry-tagged Hfq) and merged images of (a) PNPase or (b) RhlB and Hfq in 24 h N starved *E. coli* cells. (c, d) Representative fluorescence microscopy (for GFP-tagged PNPase and RhlB) and PALM (for PAmCherry-tagged Hfq) images of (c) PNPase or (d) RhlB and Hfq in 24 h N starved *E. coli* cells containing RNase E_1-700_. (e, f) Representative fluorescence microscopy (for GFP-tagged RNase E), PALM (for PAmCherry-tagged Hfq) and merged images of RNase E and Hfq in 24 h N starved *E. coli* cells devoid of (a) PNPase (Δ*pnp*) and (b) RhlB (Δ*rhlB*) cells. The arrows indicate the RNase E, PNPase, RhlB and/or Hfq foci. The %H_IM_ values are indicated in all PALM images.

### 2.5 Hfq but not H-bodies has a role in rRNA degradation in long-term N starved *E. coli*

It is well established that rRNA becomes degraded in both N and carbon (C) starved *E. coli*, and this concomitantly results in a decrease in the cellular ribosome content (Dai *et al*., 2016, Li *et al*., 2018). Consistent with this, we recently reported that 16S and 23S rRNAs are gradually degraded over the initial 24 h under N starvation, i.e., over a period of time that coincides with H-body formation (Switzer *et al*., 2020, McQuail *et al*., 2020). RNase E is an important component in the rRNA degradation pathway in *E. coli* (Sulthana *et al*., 2016). In nutrient starved *E. coli*, RNase PH initiates the degradation of 16S rRNA by removal of nucleotides from its 3’ end and this leads to the endonucleolytic cleavage of 16S rRNA by RNase E; likewise, a single endonucleolytic cleavage initiates the degradation of 23S rRNA. The resulting fragments are then degraded to mononucleotides by RNase R and RNase II. Given the role of total cellular RNA in H-body formation (Figure 3), we considered whether the regulation of rRNA during long-term N-starvation was associated with the H-body. As shown in Figure 5a, it appears that the degradation of 16S and 23S rRNAs in long-term N starved *E. coli* is dependent on Hfq as the efficiency of rRNA degradation is significantly compromised in the Δ*hfq E. coli* strain. We compared the abundance of cellular rRNA (16S and 23S rRNA) relative to the total RNA (referred to as %16S, %23S or %rRNA_TOT_; see Experimental Procedures) during exponential growth (where H-bodies do not form) and 24 h following N starvation (where H-bodies are detectable). As shown in Figure 5a and 5b, the %rRNA_TOT_ was decreased by ∼10-fold in wild-type bacteria between N+ and N-24. In contrast, %rRNA_TOT_ did not significantly differ in Δ*hfq* bacteria between N+ and N-24, suggesting that 16S and 23S rRNA are not efficiently degraded in long-term N starved *E. coli* in the absence of Hfq. Previously, we demonstrated that Hfq foci, i.e., H-bodies, do not form in *E. coli* specifically starved for C for 24 h (McQuail *et al*., 2020). Therefore, we calculated %rRNA in C starved bacteria. As shown in Figure 5a and 5b, the absence of Hfq had a significantly less pronounced affect on %rRNA at C-24 between mutant and wild-type bacteria. Therefore, we speculated whether H-bodies, which specifically form in long-term N-starved *E. coli* have any role in rRNA degradation. However, this does not seem to be the case: Firstly, RNase R which associates with Hfq for the degradation and processing of rRNA (Dos Santos *et al*., 2020) does not appear to be a constituent of the H-body (Figure 5c); secondly, the %16S and %23S rRNA between the *E. coli* strain containing RNase E_1-700_ (in which H-bodies do not form (Figure 2f) but is active for 16S and 23S rRNA degradation (Sulthana *et al*., 2016)) and the wild-type strain did not differ at N-24 (Figure 5d); thirdly, RpsB, which we used as a surrogate for 30S ribosomal protein was not present in the H-body (Figure 5e). Overall, we conclude that although Hfq has a direct role in 16S and 23S rRNA degradation in long-term N-starved *E. coli*, this process appears to happen independently of the H-body.

**Figure 5.**
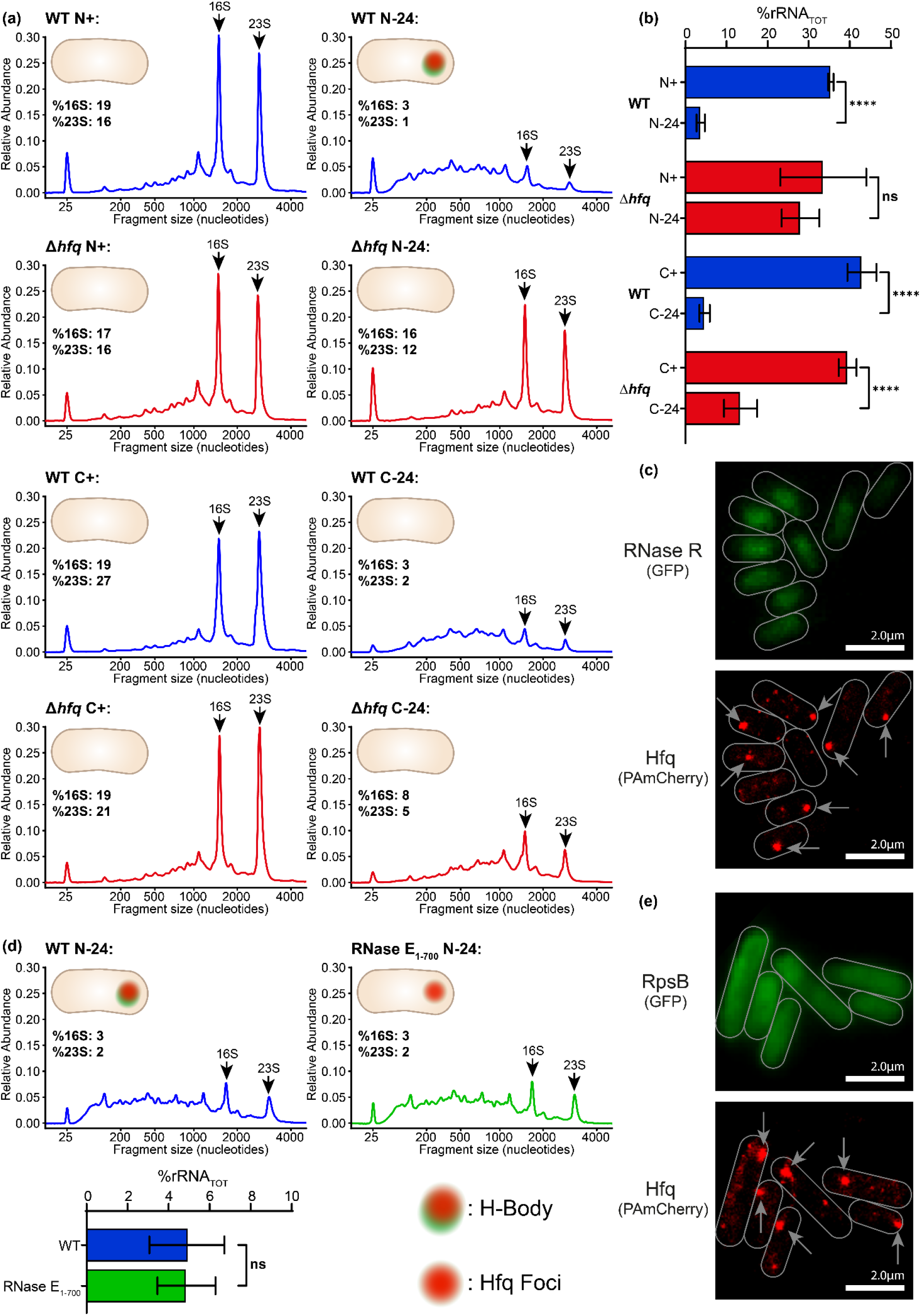
Hfq, but not H-bodies, is required for regulation of rRNA levels during long-term N-starvation. (a) Electropherograms of total RNA extracted from WT and Δ*hfq E. coli* grown under either N-limited or C-limiting conditions, sampled at the indicated time points. Graphs represent the mean RNA profile, from at least three biological replicates. The 16S and 23S rRNA peaks are indicated. The peak at 25 nucleotides is a standard marker and can be ignored. The relative abundance of 16S and 23S rRNA, as a percentage of total RNA, is indicated. (b) Graph showing relative 16S and 23S rRNA abundance (rRNA_TOT_) as a percentage of total RNA extracted from WT and Δ*hfq E. coli* at indicated time points when grown in N-limited and C-limiting conditions. Error bars represent standard deviation. Asterisks indicate significant differences between strains (****: p < 0.0001). (c) Representative fluorescence microscopy (for GFP-tagged RNase R) and PALM (for PAmCherry-tagged Hfq) images of RNase R and Hfq in 24 h N starved *E. coli* cells. (d) As in (a) with RNA extracted from 24 h N starved *E. coli* cells containing WT or RNase E_1-700_. Inset graphs show rRNA_TOT_ as a percentage of total RNA extracted from WT and bacterial containing RNase E_1-700_ at N-24. (e) Representative fluorescence microscopy (for GFP-tagged RpsB) and PALM (for PAmCherry-tagged Hfq) images of RpsB and Hfq in 24 h N starved *E. coli* cells. In (c, d), the arrows indicate the Hfq foci. In (a, e) the schematic of the *E. coli* cell indicates whether H-bodies and/or Hfq foci are present under the given condition/strain.

### 2.6 H-bodies resemble liquid-liquid phase separated subcellular assemblies

Liquid-liquid phase separation (LLPS) is a mechanism by which cells form membraneless subcompartments, a form of biomolecular condensate. LLPS occurs when proteins and nucleic acids – often associated with a common biological process – make multiple transient and multivalent interactions that causes them to condense into a dense phase that resembles liquid droplets, that coexists with a dilute phase (i.e., the cytoplasm) (Alberti *et al*., 2019). Previously Al-Husini et al showed that RNase E of *C. crescentus* forms BR-bodies by LLPS (Al-Husini *et al*., 2018). This precedence, together with our finding that the H-body is unlikely to be anchored to the inner cytoplasmic membrane (Figure 2g), prompted us to consider whether H-body formation occurs by LLPS. Therefore, as the formation of the RNase E foci (and by extension the formation of PNPase and RhlB foci) are dependent of the formation of the Hfq foci (Figure 2e), we used the Hfq foci as a surrogate to determine whether H-bodies form by LLPS. Sensitivity to hexanediol (HEX) - an aliphatic alcohol that disrupts subcellular assemblies that form by LLPS - is widely utilised as a useful, albeit not definitive, indicator of LLPS in bacterial cells (Al-Husini *et al*., 2018, Ladouceur *et al*., 2020b). Therefore, we treated *E. coli* cells from N-24, in which Hfq foci had formed, with HEX for 30 min. As shown in Figure 6a, the Hfq foci dispersed following the addition of HEX. Further, the Hfq foci reassembled when the HEX was washed out of the culture; although, we note that some cells did not contain any or contained two Hfq foci following washing out HEX (Figure 6a). Consistent with the observation that the formation of the RNase E foci is dependent on the Hfq foci, timelapse PALM analysis revealed that H-bodies also dispersed following HEX treatment (Figure 6b). However, as sensitivity to HEX is not a definitive test for LLPS (Alberti *et al*., 2019), we considered whether the Hfq foci (and thus H-bodies) conform with other known properties of LLPS condensates. A comprehensive review by Azaldegui et al (Azaldegui *et al*., 2021), proposed the following criteria for determining whether a structure is formed by LLPS: *(i)* condensates are spherical in nature due to surface tension, *(ii)* condensate droplets can fuse upon contact, *(iii)* molecules within condensates remain mobile and *(iv)* are able to diffuse across the condensate boundary, and *(v)* condensate formation occurs in a concentration dependent manner. It seems that the Hfq foci meet three (*i, iii* and *iv*) of these five criteria: As seen in figures 1-4, the Hfq foci, i.e., the H-body, is consistently spherical-shaped in 2D space and are therefore likely to be spherical in 3D space. The tracking of individual molecules of Hfq is possible (McQuail *et al*., 2020) and analysis of individual trajectories of Hfq molecules in 24 h N starved *E*.*coli* reveals that individual Hfq molecules are not only mobile *within* the Hfq foci (Figure 6c, trajectories A,B), but that individual Hfq molecules are able to diffuse *across* the Hfq foci boundary (Figure 6c, trajectories C, D). However, given that only a single Hfq-foci is ever present within the cell, we are unable to determine whether these condensates are able to fuse (criteria *ii*). It is possible that smaller clusters of Hfq fuse but, as the Hfq-foci form over the course of several hours (McQuail *et al*., 2020), it is not technically possible to perform time-lapse microscopy on cells whilst maintaining the conditions required for Hfq-foci formation. Further, we are currently unable to demonstrate the concentration dependence for Hfq-foci formation (criteria *v*) as the complete composition of the H-body is not known, which hinders *in vitro* analysis at this time. Overall, based on the fact that the Hfq foci are sensitive to HEX and meet criteria *i, iii* and *iv* of LLPS condensates, we conclude that the Hfq foci, i.e., H-bodies, in long-term N starved *E. coli* cells physically resemble subcellular assemblies that form by LLPS.

**Figure 6.**
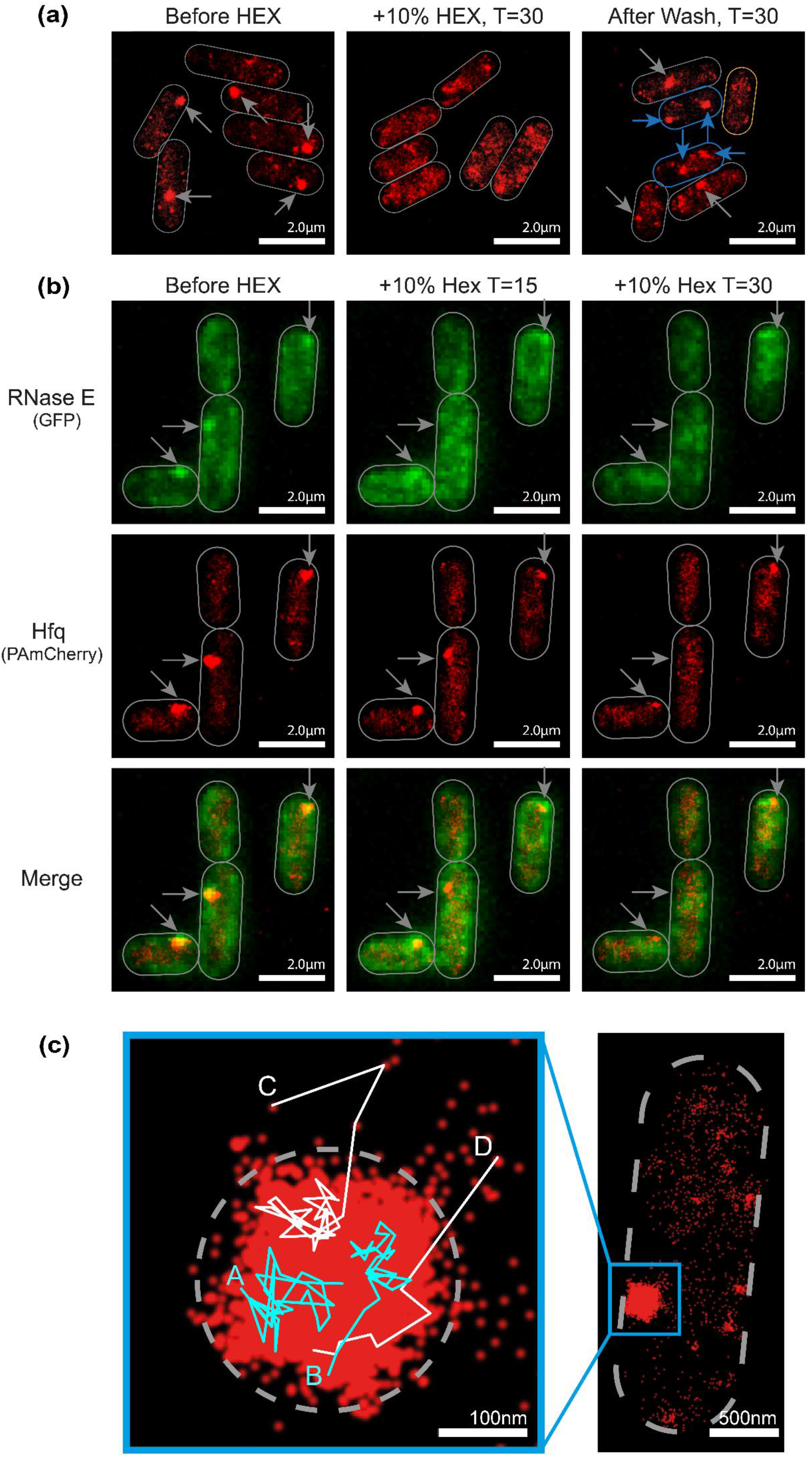
H-bodies resemble liquid-liquid phase separated subcellular assemblies. (a) Representative PALM images of PAmCherry-tagged Hfq in 24 h N starved *E. coli* cells before (left), after (middle) 30 min treatment with 10% (v/v) hexanediol and 30 min after removal of hexanediol from the culture (right). Cells that contain no Hfq foci or contain two Hfq foci following HEX wash out are showin in yellow and blue, respectively. (b) Representative images from timelapse fluorescence microscopy (for GFP-tagged RNase E), PALM (for PAmCherry-tagged Hfq) and merged images of RNase E and Hfq in 24 h N starved *E. coli* cells before and after 15 min and 30 min treatment with 10% (v/v) hexanediol. The arrows indicate the RNase E and/or Hfq foci. (c) Hfq remains mobile within the Hfq foci and can diffuse across the Hfq foci boundary. Representative individual trajectories of Hfq within the Hfq foci. Trajectories A and B show movement of Hfq *within* the Hfq foci; trajectories C and D show movement of Hfq *across* the Hfq foci boundary.

## 3. Discussion

The importance of the association between RNase E and Hfq to the post-transcriptional fate of RNA in bacteria is undisputed (Mackie, 2013, Vargas-Blanco & Shell, 2020, Morita & Aiba, 2011, Ikeda *et al*., 2011, Morita *et al*., 2005). We recently reported that Hfq molecules assemble into a large focus-like structure in long-term N starved *E. coli*. Although we demonstrated that the Hfq foci are important for *E. coli* to adapt to long-term N starvation (McQuail *et al*., 2020), their composition, mechansims of function and prevalence remain elusive. In this study, we focused on the composition of the Hfq foci. Prompted by the established associations between Hfq/sRNA and the RNA degradosome in *E. coli* (Bruce *et al*., 2018, Morita *et al*., 2005) and the recent discovery that the RNA degradosome can form foci-like assemblies in *C. crescentus* (Al-Husini *et al*., 2018, Al-Husini *et al*., 2020), we focused our analysis on investigating whether the Hfq foci contained components of the RNA degradosome. Indeed, we have demonstrated that RNase E, PNPase and RhlB, which constitute the minimal RNA degradosome in *E. coli*, colocalise with the Hfq foci to form a structure which we term the H body. Further, to the best of our knowledge, this is the first time that Hfq and the RNA degradosome have been visualised together in *live* bacteria. As RNase E and Hfq colocalise even in mutant bacteria devoid of either PNPase or RhlB, it is plausible, but impossible to decipher from our experiments, that H-bodies consists of (at least) two mutually exclusive multiprotein complexes, namely, Hfq/sRNA and the minimal RNA degradosome and Hfq/sRNA and RNase E. The existence of the latter complex is supported by co-immunoprecipitation experiments conducted by Morita et al in *E. coli* cells devoid of either PNPase or RhlB (Morita *et al*., 2005).

In recent elegant studies by the Arraiano group, Hfq has been demonstrated to contribute to the biogenesis (Andrade *et al*., 2018) and processing and degradation (Dos Santos *et al*., 2020) of rRNA in *E. coli*. Notably, our study now extends this observation and demonstrates a role for Hfq in rRNA degradation in long-term N (and less so in C) starved *E. coli* that occurs independently of the H-bodies. We thus speculate that the in the absence of Hfq, sustained biosynthetic activity in N starved bacteria cells would aberrantly deplete cellular resources, which, compounded by their inability to form H-bodies would ultimately compromise viability.

Remarkably, the size (∼0.3 µm in diameter), behaviour (dispersion when stress is alleviated), longevity and physical properties (formation by LLPS like mechanism) of the H-bodies bear similarities to the BR-bodies seen in α-proteobacteria (Al-Husini *et al*., 2018, Al-Husini *et al*., 2020) and analogous assemblies in stressed eukaryotic cells, such as P bodies or stress granules (Luo *et al*., 2018). As P-bodies and stress granules are sites of RNA storage and turnover in stressed eukaryotic cells, and based on the observation that the depletion of total cellular RNA affects H-body formation (Figure 3), we speculate that RNA is a constituent of the H-bodies. As previously reported by us, due to the poor permeability of bacteria at N-24, attempts to stain the H-bodies with RNA specific stains were unsuccessful (McQuail *et al*., 2020) and experiments are currently underway to understand the nucleoprotein composition of H-bodies at the genome-wide scale.

Membraneless compartmentalisation of RNA associated processes by LLPS is emerging to be pervasive in bacteria and, in addition to BR-bodies in α-proteobacteria, the RNA polymerase has also been shown to undergo LLPS in *E. coli* (Ladouceur *et al*., 2020a). Our results suggest that the H-bodies represent a ‘membraneless compartment’ determining the post-transcriptional fate of RNA in long-term N starved *E. coli* cells. The fact that H-bodies disperse when the N starvation stress is alleviated (Figure 2d) suggests that H-bodies contribute to the adaptive response of *E. coli* to long-term N starvation. However, a limitation of our study is that it is impossible to directly establish whether H-bodies are required for optimal viability in long-term N starved *E. coli* due to the pleiotropic roles H-body associated proteins (i.e., Hfq, RNase E, PNPase and RhlB) play in gene regulation and bacterial stress response. Nevertheless, using the RNase E_1-700_ variant (that does not form H-bodies; Figure 2f), which is perhaps the most conservative mutant to use, we can demonstrate that the H-bodies could indeed contribute to optimal viability of *E. coli* experiencing long-term N starvation (Supplementary Figure 3).

Interestingly, intrinsically disordered regions of RNA binding proteins are often associated with LLPS in both eukaryotic and prokaryotic systems. In fact, the RNase E of *C. crescentus* requires its intrinsically disordered carboxyl terminal region to undergo LLPS and form BR-bodies (Al-Husini *et al*., 2018). Our data demonstrates that Hfq foci formation is a requirement for RNase E foci to form and colocalise with the Hfq foci, i.e., formation of H-bodies. Intriguingly, the intrinsically disordered carboxyl terminal region of Hfq is not required for Hfq foci formation (McQuail *et al*., 2020). However, RNase E_1-700_, which lacks its intrinsically disordered region, is unable to form foci and colocalise with the Hfq foci. This could suggest that the intrinsically disordered region of RNase E might be required for foci formation by LLPS and association with the Hfq foci. However, given that associaton with Hfq is a requirement of RNase to form foci, we cannot exclude that the inability of RNase E to interact with Hfq (because it is lacking the carboxyl terminal region), rather than the absence of the intrinsically disordered region *per se*, is the contributing factor for the inability of RNase E_1-700_ to form the foci. Therefore, it is possible that H-body formation by LLPS and the physically properties of H-bodies are influenced by other, yet to be identified, interacting partners.

Overall, although many fundamental properties of H-bodies in bacterial stress response remain to be determined, this study has uncovered that in long-term N starved *E. coli*, the association with the RNA degradosome via a LLPS like mechanism and rRNA degradation are hitherto unreported and mutually exclusive properties of the ubiquitous bacterial RNA chaperone Hfq.

## 4. Experimental Procedures

### Bacterial strains and plasmids

All strains used in this study were derived from *Escherichia coli* K-12 and are listed in Supplementary Table 1. The Hfq-PAmCherry was made as previously described (McQuail *et al*., 2020). Tagged genes (*rne*-msfGFP, *rne*(Δ565−585)-eGFP, *rne*700-msfGFP, *pnp*-msfGFP, *rhlB*-msfGPF and *rnr*-msfGFP) and gene deletions (Δ*hfq*, Δ*pnp* and Δ*rhlB*) were introduced into the *E. coli* strain containing Hfq-PAmCherry and derivative strains as described previously (Brown *et al*., 2014). Briefly the tagged or knockout alleles were transduced using the P1vir bacteriophage from the appropriate donor strains listed in Table 1. The *rne*-msfGFP strain (SAJ232) and *rne700*-msfGFP strains (SAJ233) were constructed as described previously (Strahl *et al*., 2015).

### Bacterial growth conditions

Bacteria were grown in Gutnick minimal medium (33.8 mM KH_2_PO_4_, 77.5 mM K_2_HPO_4_, 5.74 mM K_2_SO_4_, 0.41 mM MgSO_4_) supplemented with Ho-LE trace elements (Atlas, 2010), 0.4 % (w/v) glucose as the sole C source and NH_4_Cl as the sole N source. Overnight cultures were grown at 37 °C, 180 r.p.m. in Gutnick minimal medium containing 10 mM NH_4_Cl. For the N starvation experiments, 3 mM NH_4_Cl was used. The proportion of viable cells in the bacterial population was determined by measuring CFU/ml from serial dilutions on lysogeny broth agar plates. To observe Hfq and RNase E foci dissipation, 25 ml of the 24 h N starved culture was centrifuged at 3,200 *g* and resuspend in fresh Gutnick minimal medium containing 0.4% (w/v) glucose and 3 mM NH_4_Cl and grown at 37 °C, 180 r.p.m. for 3 hours. For carbon starvation experiments, bacteria were grown in Gutnick minimal medium containing 22 mM glucose and 10 mM NH_4_Cl as previously described (McQuail *et al*., 2020).

### Microscopy

For the PALM experiments, the bacterial cultures were grown as described above and samples were taken at the indicated time points, then imaged and analysed as previously described (Stracy *et al*., 2015, Endesfelder *et al*., 2013). Briefly, 1 ml of culture was centrifuged, washed and resuspended in a small amount of Gutnick minimal medium without any NH_4_Cl; samples taken at N+ were resuspended in Gutnick minimal medium with 3 mM NH_4_Cl. One microlitre of the resuspended culture was then placed on a Gutnick minimal medium agarose pad (1x Gutnick minimal medium without any NH_4_Cl with 1% (w/v) agarose); samples taken at N+ were placed on a pad made with Gutnick minimal medium with 3 mM NH_4_Cl. Cells were imaged on a PALM-optimized Nanoimager (Oxford Nanoimaging, www.oxfordni.com) with 15 millisecond exposures, at 66 frames per second over 10,000 frames. Photoactivatable molecules were activated using 405 nm and 561 nm lasers. GFP images were acquired as above, except using only a 488 nm laser, with single, 15-100 millisecond exposures, depending on GFP levels within the cell. Typically, a field-of-view typically consisting of ∼100-200 bacterial cells.

### Total RNA analysis

Cultures were grown as described above and sampled at the following time points: during exponential growth (N+, C+) and 24 h under starvation (N-24, C-24), during N and C starvation respectively. At least three biological replicates were taken at each time point. Cell samples were acquired and stabilized at specified time points using Qiagen RNA Protect reagent (76526; Qiagen). RNA extractions were performed using the PureLink RNA minikit (12183025; Invitrogen) and DNase set (12185010; Invitrogen) according to the manufacturers’ protocols. Electropherograms of total RNA integrity were generated using an Agilent Bioanalyzer 2100 instrument (G2939BA; Agilent) and RNA 6000 Nano kit (5067-1511; Agilent) per the manufacturer’s protocol. The relative abundance of 16S and 23S rRNA of RNA samples was determined by the Agilent Bioanalyzer software suite.

## Acknowledgements

We thank Lynn Burchell and Leonora Poljak who helped produce some of data and *E. coli* strains. This work was supported by grants from the Wellcome Trust (100958), Leverhulme Trust (RPG-2020-050) and BBSRC (BB/V000284/1) to SW. We thank Amy Switzer and Aline Tabib-Salazar for comments on the manuscript.

## Author Contributions

Experimental design and planning (JM and SW), production and analysis of data (JM and SW), preparation of the manuscript (SW, JM, AJC).

---

**Supplementary Figure 1.**
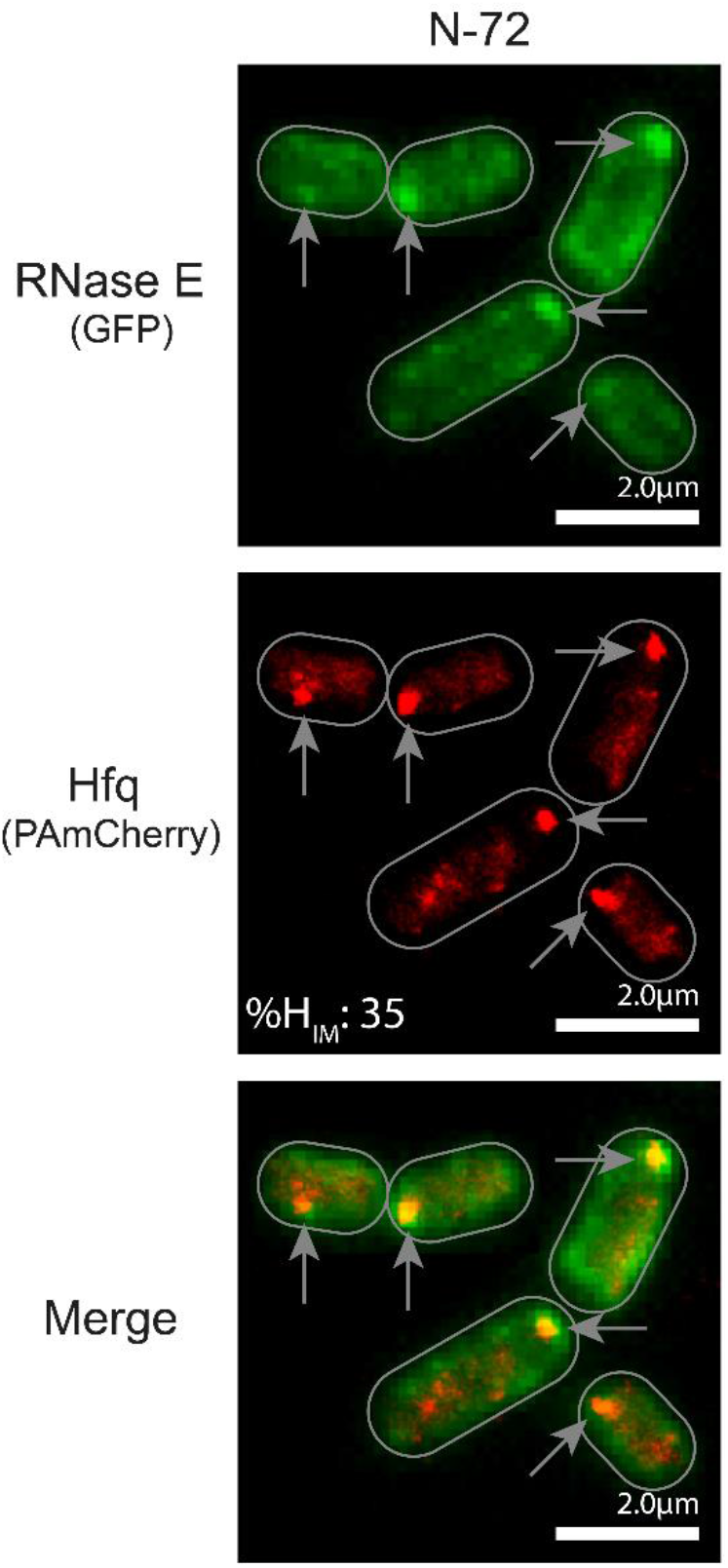
The RNase E foci persist for up to 72 h. Representative fluorescence microscopy (for RNase E), PALM (for Hfq) and merged images of RNase E and Hfq in 72 h N starved *E. coli* cells.

**Supplementary Figure 2.**
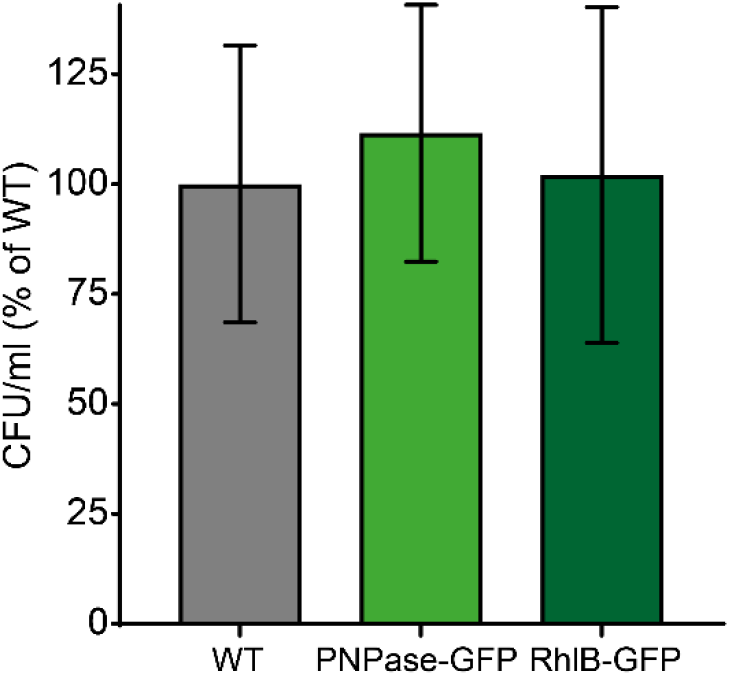
The GFP tags on PNPase and RhlB do not affect cell viability during long-term N-starvation. Graph showing the proportion of viable cells in cultures of 24 h N starved WT and GFP-tagged PNPase and RhlB-GFP containing *E. coli*, measured by counting CFU and shown as a percentage of WT. Error bars represent standard deviation (n = 3).

**Supplementary Figure 3.**
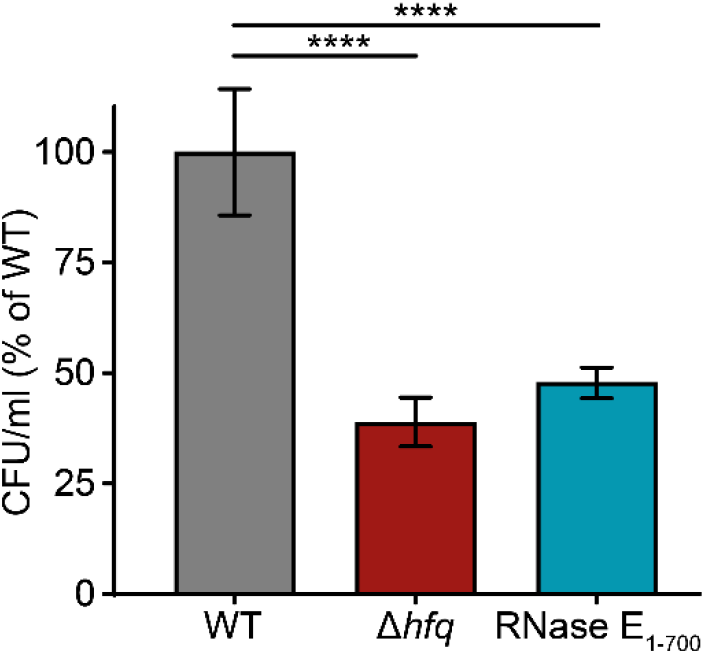
Graph showing the proportion of viable cells in cultures of 24 h N starved WT, Δ*hfq* and *rne*_1-700_ *E. coli* measured by counting CFU and shown as a percentage of WT. Error bars represent standard deviation (n = 3). Asterisks indicate significant differences between strains (****: p < 0.0001).

**Supplementary Table 1.** -*E. coli* strains and plasmids used in this study

| **Strains** |  |  |
| --- | --- | --- |
| Name | Description | Source or Reference |
| Wild-type MG1655 | *E. coli* K-12 *rph-1* | E. coli Genetic Stock Centre |
| Wild-type (BW25113) | *E. coli* K-12 (*araD-araB*)567 $\Delta$(*rhaD-rhaB*) 568 Δ*lacZ*4787 (::rrnB-3) *hsdR*514 *rph-1* | E. coli Genetic Stock Centre |
| Rne-msfGFP (SAJ232) | NCM3416 *rne*(1-1061)-*msfGFP-cat* | This study |
| RneΔMTS-eGFP (Kti491) | NCM3416 *rne*($\Delta$ 565-585)-*efGFP-cat* | [1] |
| Hfq-PAmCherry | MG1655 *hfq-PAmCherry-kan* | [2] |
| Rne-msfGFP Hfq-PAmCherry | MG1655 *rne*(1-1061)-*msfGFP-cat hfq-PAmCherry-kan* | This Study |
| RneΔ565-585-eGFP Hfq-PAmCherry | MG1655 *rne*(Δ565-585)-*efGFP -cat hfq-PAmCherry-kan* | This Study |
| Δ*hfq* (JW4130) | BW25113 Δ*hfq::kan* | E. coli Genetic Stock Centre |
| Δ*hfq* Rne-msfGFP | MG1655 Δ*hfq::kan rne*(1-1061)-*msfGFP-cat* | This Study |
| Rne700-msfGFP (SAJ233) | NCM3416 *rne*(1-700)-*msfGFP-cat* | This study |
| Rne700-msfGFP Hfq-PAmCherry | MG1655 *rne*(1-700)-*msfGFP-cat hfq-PAmCherry-kan* | This Study |
| Δ*pnp* (JW5851) | BW25113 Δ*pnp::kan* | E. coli Genetic Stock Centre |
| Δ*rhlB* (JW3753) | BW25113 Δ*rhlB::kan* | E. coli Genetic Stock Centre |
| Δ*pnp* Rne-msfGFP Hfq-PAmCherry | MG1655 *rne*(1-1061)-*msfGFP-cat hfq-PAmCherry Δpnp::kan* | This Study |
| Δ*rhlB* Rne-msfGFP Hfq-PAmCherry | MG1655 *rne*(1-1061)-*msfGFP-cat hfq-PAmCherry ΔrhlB::kan* | This Study |
| Pnp-msfGFP (SLM013) | NCM3416 *pnp-msfGFP-cat* | Gift from Lina Hamouche |
| RhlB-msfGFP (SLM021) | NCM3416 *rhlB-msfGFP-cat* | Gift from Lina Hamouche |
| Pnp-msfGFP Hfq-PAmCherry | MG1655 *pnp-msfGFP-cat hfq-PAmCherry-kan* | This Study |
| RhlB-msfGFP Hfq-PAmCherry | MG1655 *rhlB-msfGFP-cat hfq-PAmCherry-kan* | This Study |
| Rne701-FLAG (TM528) | W3110mic *rne*(1-701)-*Flag-cat* | [3] |
| Rne701-FLAG Pnp-msfGFP Hfq-PAmCherry | MG1655 *rne*(1-701)-*Flag-cat pnp-msfGFP hfq-PAmCherry-kan* | This Study |
| Rne701-FLAG RhlB-msfGFP Hfq-PAmCherry | MG1655 *rne*(1-701)-*Flag-cat rhlB-msfGFP hfq-PAmCherry-kan* | This Study |
| Rnr-msfGFP Hfq-PAmCherry | MG1655 *rnr-msfGFP-cat hfq-PAmCherry-kan* | This Study |
| RpsB-msfGFP Hfq-PAmCherry | MG1655 *rpsB-msfGFP-cat hfq-PAmCherry-kan* | This Study |
| **Plasmids** |  |  |
| Name | Description | Source or Reference |
| pBAD24-*hfq*-FLAG | pBAD24 expressing *hfq*-3xFLAG under an arabinose inducible promoter | Gift from Jörg Vogel, |

